# Identification and visualisation of differential isoform expression in RNA-seq time series

**DOI:** 10.1101/155135

**Authors:** María José Nueda, Jordi Martorell-Marugan, Cristina Martí, Sonia Tarazona, Ana Conesa

## Abstract

As sequencing technologies improve their capacity to detect distinct transcripts of the same gene and to address complex experimental designs such as longitudinal studies, there is a need to develop statistical methods for the analysis of isoform expression changes in time series data. Iso-maSigPro is a new functionality of the R package maSigPro for transcriptomics time series data analysis. Iso-maSigPro identifies genes with a differential isoform usage across time. The package also includes new clustering and visualization functions that allow grouping of genes with similar expression patterns at the isoform level, as well as those genes with a shift in major expressed isoform. The package is freely available under the LGPL license from the Bioconductor web site (http://bioconductor.org).

## 1 Introduction

Alternative splicing (AS) is a common mechanism of higher eukaryotes to expand transcriptome complexity and functional diversity. The expression of alternative isoforms of many genes is a developmentally regulated process (Vuong *et al*., 2016) and AS has been shown to occur as response to environmental cues (AlShareef *et al*., 2017). Hence, there is an interest in studying the dynamics of AS. Deep sequencing methods currently used in transcriptomics research allow for the study of AS. Reads that map to splice junctions can be used to estimate splicing events, while several transcript recontruction and quantification methods have been published that enable inference of isoform expression with different levels of accuracy (Steijger *et al*., 2013). More recently, long-read sequencing platforms, which allow transcript identification without the need of a reconstruction step, have become available to boost the study of isoform regulation (Au *et al*., 2013). While many algorithms have been developed for differential AS analysis most of these approaches target pair-wise comparisons (i.e. Andres *et al*., 2012 and Matthew *et al*., 2015) and have not yet developed specific models that integrate time course with differential splicing analysis. The DICESeq method was designed for a better estimation of isoform expression in time series, but does not implement a specific strategy for obtaining differentially expressed isoforms (Huang and Sanguinetti, 2016). Topa and Honkela proposed to model time series as a Gaussuian process and isoform levels as proportions over the total gene expression to then evaluate splicing as the change between time-dependent and time-independent models (Topa and Honkela, 2016). This approach requires large datasets to fit parameters and does not consider comparisons between multiple series, such as treatment and control. Compared to genes or transcripts, the analysis of differential isoform expression in time course experiments poses a number of specific challenges. Different transcripts of the same gene may vary in their time trajectories and the analysis algorithm should be able to identify those genes where isoform profiles change differently in a significant manner, i.e. those genes that are differentially spliced across time. Additionally, joint visualisation of significant splicing changes is complicated by the fact that genes have different number of isoforms and hence data does not fit into the structure of traditional clustering, where the same number of data points is required for each feature. Therefore, novel clustering strategies should be envisioned to group genes expressing their isoforms in a similar fashion. Finally, transcripts of the same gene have frequently very different expression levels, with one “major” isoform bearing most of the expression signal and alternative isoforms being lowerly expressed. Ideally, analysis approaches should be able to account for this characteristic and identify those cases where genes change their major isoform in the course of time.

maSigPro is an R package specifically designed for the analysis of multiple time course transcriptomics data. maSigPro fits an optimized polynomial linear model to describe the dynamics of gene expression in one or multiple experimental conditions and selects genes with significant model coefficients (Conesa *et al*., 2006). The package incorporates a clustering function to visualize genes with similar profiles. maSigPro was initially developed for microarrays and later updated to model count data (Nueda *et al*., 2014). In this paper we present Iso-maSigPro, a further adaptation to study differential isoform usage in time course RNA-seq experiments. We implement a new function to model differential splicing and combine this with differential transcript expression analysis to identify genes where isoforms change expression across time. Novel query and visualisation functions allow selecting genes with the strongest isoform switches and grouping genes with similar time-dependent AS patterns.

## 2. Methods

### 2.1 Model

The Generalized Linear Model (GLM) described in Nueda *et al*., 2014 to study the gene expression value *y_i_* at observation *i*, time *t_i_* and *s* experimental conditions (i.e, treatments, tissues, strains, etc) identified by s — 1 binary variables 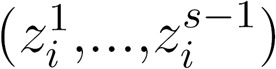 can be written as follows (when considering s = 2 and linear effects):

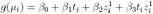

Being *μ_i_ = E(y_i_)*, *g* a monotonic differentiable function called ‘link function’, which characterizes the GLM model, and *β_0_, β_1_, β_2_, β_3_* the coefficients to estimate.

#### 2.1.1 Model for Differentially Spliced Genes (DSG) across time

For each multi-isoform gene two models are created, identifying *J* isoforms with *J —* 1 binary variables (*I^1^,…,I^J-1^*). The reference model, *M*_0_, considers there exist only constant differences between isoforms and the global gene model, *M*_1_, considers the possibility of a time vs condition vs isoform interaction. *M_0_* imposes parallel profiles to the different isoforms, in contrast *M*_1_ allows modeling different profiles and hence captures the differential splicing cases. For instance, for a gene with two isoforms, two experimental conditions or series and linear effects:

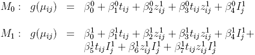

Being *μ_ij_ = E(yij)* the expected value for observation *i* and isoform *j*.

To evaluate the statistical significance of the interaction, both models are compared for each gene. In GLMs hypothesis testing is based on the log-likelihood ratio statistic (McCullagh and Nelder, 1989; Wood, 2006).

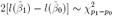

Where 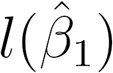 is the maximized likelihood of the complete model with 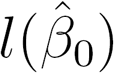 the likelihood of the reference model with *p_0_* parameters, being *p_0_ < p_1_*.

#### 2.1.2 The Iso-maSigPro functions

Seven new functions have been added to the maSigPro package to enable analysis of differentially expressed isoforms. Figure 1 shows the analysis pipeline and novel Iso-maSigPro functions:

1. *IsoModel()* implements the DS models *M_0_* and *M_1_* for each multi-isoform gene, using the polynomial model obtained with the generic *make, design, matrix()* maSigPro function that best describes the experimental design. The comparison of both models gives as a result a FDR-corrected p-value of differential splicing.
2. Transcripts from significant DSGs are then subjected to regular Next-maSigPro analysis to detect Differentially Expressed Transcripts (DETs).
3. *IsoModel()* returns a list of DSGs together with the estimated models of associated isoforms to be used as input in *getDS()* function to obtain a selection of DSGs at a preestablished level of goodness of fit for each model.
4. Downstream analysis can be performed with functions *seeDS(), tableDSf), getDSPat-tem(), PodiumChange()* and *IsoPlot()*, that cluster, select and visualize patterns of isoform change.

Note that in this formulation, it is possible that a gene is called DSG but no significant DETs of that gene are found under the significance level, goodness of fit and multiple testing correction constraints of the regular maSigPro analysis.

**Figure 1:**
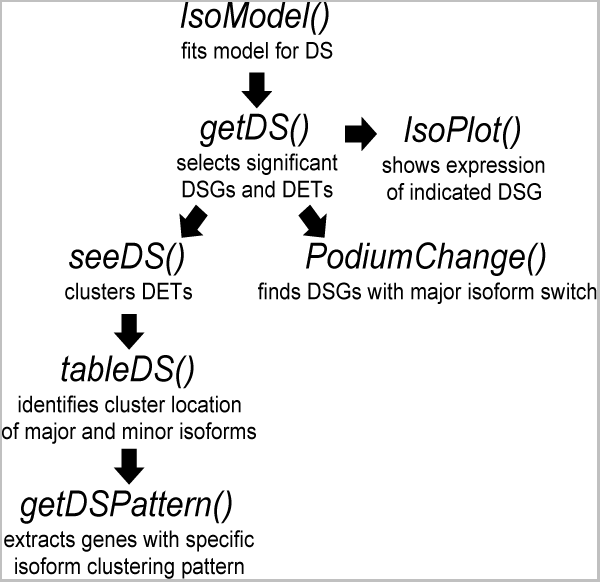
Workflow for Iso-maSigPro analysis.

### 2.2. Visualization

Typically, maSigPro will cluster features according to their expression pattern in all experimental conditions. This option is still available for all differential transcripts regardless of their parent gene. The Iso-maSigPro framework allows for two additional visualisation functions to study differential splicing results: differential splicing clustering and major isoform switch.

#### 2.2.1 Differential splicing clustering with *seeDS()* and *tableDS()*

The clustering strategy implemented with these two functions aims to identify groups of DSGs with similar isoform expression patterns. First, *seeDS()* takes DETs-either from DSGs or in combination with DETs from single-transcript genes-and clusters them into *k* groups with any of the available maSigPro clustering approaches to define transcriptional patterns globally present in the data. Next, *tableDS()* identifies, for each DSG, the cluster(s) their DETs belong to and labels gene transcripts as major (here defined as the isoform with the highest total expression across conditions) or minor isoforms. This information is used to create a classification table that indicates the distribution of DETs of DSG across different clusters. By evaluating the classification table with the cluster profiles, the user can readily identify genes with strong or subtle expression differences among their set of isoforms.

#### 2.2.2 PodiumChange()

This function returns DSGs that undergo a switch of their most expressed isoform during the time course. *PodiumChange()* can be applied taking into consideration only DETs or all isoforms of DSGs. This last option is interesting when the DSG has only one isoform called as DET. The function takes as input the result of *getDS()* and returns a list of genes with podium changes. The function can detect changes at any time point (eventual changes), for an indicated experimental condition or at specific subranges of time and experimental conditions. Finally an isoform-resolved expression profile graph of genes with podium changes can be plotted with the *IsoPlot()* function to reveal the switch among isoforms.

#### 2.2.3 IsoPlot()

This function provides gene-level plots of the expression profiles of all transcripts in the input genes. Optionally, the user can choose to visualize all transcripts or only DETs of the selected genes. Typically, *IsoPlot()* will be used to inspect specific genes identified by the *PodiumChange()* or the *tableDS()* functions.

## 3 Results

The described methodology has been applied to the analysis of a published RNA-seq dataset (GEO accession GSE75417) describing a mouse B-cell differentiation course from the pre-BI (cycling or Hardy fraction C’) stage to the pre-BII (or Hardy fraction D) stage, where B-cell progenitors undergo growth arrest and differentiation. The process is triggered by the induction of the expression of the B-cell specific transcription factor Ikaros. Transcripts were quantified with eXpress (Roberts and Pachter, 2013). A total of 34,156 transcripts belonging to 12,572 genes are available, of which 6,882 genes expressed more than one transcript and 28,466 transcripts belong to multi-isoform genes. Data consist of 6 time points (0, 2, 6, 12, 18 and 24 hours after Ikaros induction), two experimental conditions (Control and Ikaros-induced) and three biological replicates per time and experimental condition.

### 3.1 Identification of DSGs

The *IsoModel()* function gave as overall result the selection of 347 DSGs containing a total of 1,239 transcripts. Of these, 665 also had significant time course changes (DETs). For 37 genes, we could not find individual differentially expressed transcripts, suggesting these genes had subtle expression changes that could be detected only when different isoforms profiles were compared. Table 1 summarizes results.

**Table 1:** IsoModel results on the B-cell data. Number of DSGs and DETs in differentially spliced genes.

| #DETs by DSGs | 0 | 1 | 2 | 3 | 4 | 5 | 6 | 8 | 9 | 10 | Total |
| --- | --- | --- | --- | --- | --- | --- | --- | --- | --- | --- | --- |
| #DSGs | 37 | 129 | 103 | 30 | 22 | 16 | 6 | 1 | 2 | 1 | **347** |
| #DETs | 0 | 129 | 206 | 90 | 88 | 80 | 36 | 8 | 18 | 10 | **665** |

### 3.2 Clustering of DSGs

We applied the function *seeDS()* to group our 665 DETs into 6 clusters (Figure 2) and obtained the cluster assignment of their major and minor significant isoforms (Table 2 and Supplementary tables).

**Table 2:** *tableDS()* results. Number of DSGs with major and minor isoforms in *seeDS()* clusters.

| Cluster of minor isoform(s) | Cluster of major isoform |  |  |  |  |  |
| --- | --- | --- | --- | --- | --- | --- |
|  | 1 | 2 | 3 | 4 | 5 | 6 |
| 1 | 24 | 6 | 7 | 8 | 0 | 2 |
| 1&2 | 13 | 7 | 1 | 1 | 0 | 0 |
| 1&2&3&4 | 0 | 0 | 1 | 0 | 0 | 0 |
| 1&2&4 | 0 | 1 | 0 | 1 | 0 | 0 |
| 1&3 | 0 | 0 | 0 | 1 | 0 | 0 |
| 1&4 | 1 | 0 | 1 | 1 | 0 | 0 |
| 1&5 | 0 | 1 | 0 | 0 | 2 | 0 |
| 1&6 | 3 | 1 | 1 | 0 | 0 | 1 |
| 2 | 15 | 13 | 2 | 1 | 0 | 0 |
| 2&3&4 | 0 | 0 | 1 | 0 | 0 | 0 |
| 2&6 | 0 | 1 | 0 | 0 | 0 | 0 |
| 3 | 4 | 0 | 15 | 10 | 0 | 0 |
| 3&4 | 0 | 0 | 6 | 4 | 0 | 0 |
| 3&6 | 0 | 0 | 0 | 0 | 0 | 1 |
| 4 | 5 | 0 | 5 | 5 | 0 | 0 |
| 5 | 1 | 0 | 0 | 0 | 1 | 0 |
| 6 | 2 | 0 | 1 | 1 | 0 | 2 |
|  | 68 | 30 | 41 | 33 | 3 | 6 |

**Figure 2:**
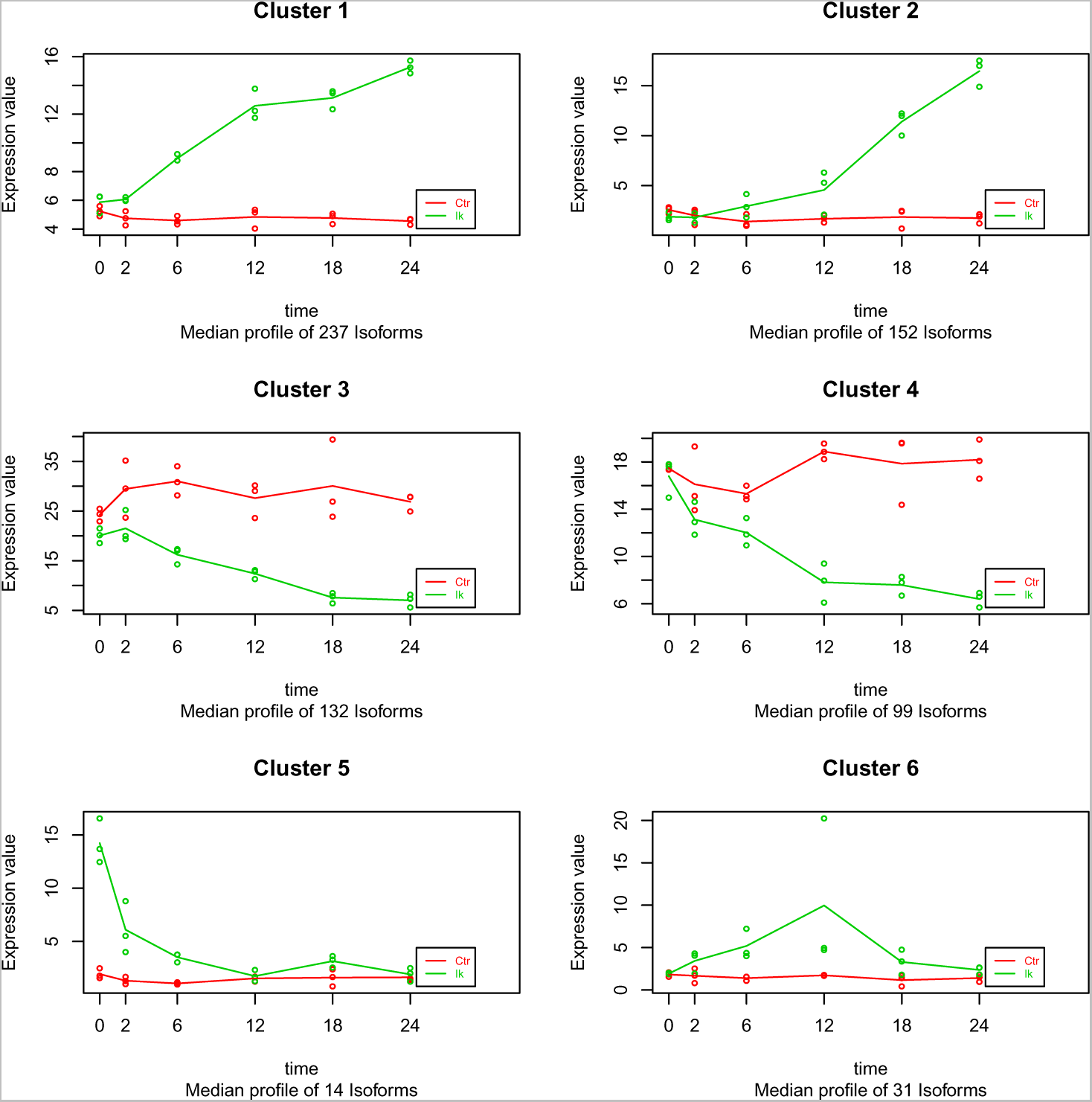
Output of *seeDS()*. Clusters of 665 DETs belonging to 347 DSGs.

We observed that, in most cases, major and minor isoforms of the same gene follow the same transcriptional pattern. For example, out of the 68 genes with major isoforms in cluster 1 (upregulation in the Ikaros series), 24 cases had minor isoforms belonging to the same cluster. For other 15 genes, the secondary transcripts fall into cluster 2, which also represents upregulation in Ikaros but with a time delay. Finally, 13 of these 68 genes have their minor forms spread between clusters 1 and 2. For a small number of other genes, expression of major and minor isoforms followed very different trajectories. For example, 7 genes had major isoforms in cluster 3 (downregulation in Ikaros) but secondary isoforms in cluster 1 (Ikaros upregulation). Figure 3A shows an example of one such gene. The transcription factor *Nfkb2* is expressed in our system with two isoforms with opposite expression profiles. Isoform ENSMUST00000073116 is expressed in the pre-BI stage and drops sharphly after Ikaros induction, while isoform ENSMUST00000011881 increased levels after 6 h. Both isoforms code for the same protein but have different 5'UTR regions that contain distinct regulatory signals (Supplementary figure S1).

**Figure 3:**
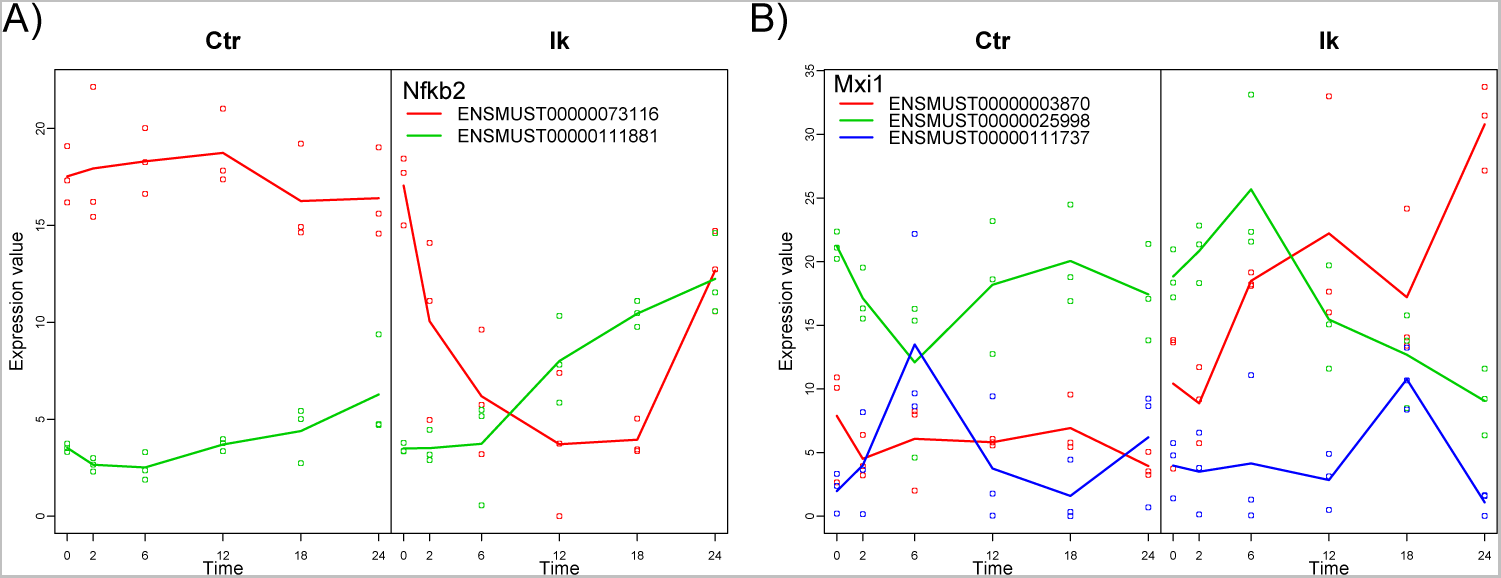
IsoPlotQ examples. A) Nfkb2 gene has isoforms in cluster 1 and 4. B) Mxi1 is a podium change gene.

### 3.3 Identification of genes with a switch of major isoform

We applied the function *PodiumChange()* to the 347 DSGs and identified 127 cases where a change in the most expressed isoform was present at any time. To select genes where the major isoform switch could be associated to the transition from cycling to differentiation stages in B-cell development, we used the *time.points* parameter in *PodiumChange()* to require a change in the last 2 time points, which resulted in 37 genes (Supplementary tables). Figure 3B shows an example of one such gene (*Mxi1*) plotted with the function *IsoPlot()*. Mxi1 is a transcriptional repressor and an antagonist of Myc, a key transcription factor that regulates B-cell differentiation and is downregulated after Ikaros induction (Ma *et al*, 2010). Interestingly, *Mxi1* changes its most expressed isoform from Mxi1-202 (ENSMUST0000025998) to Mxi1-201 (ENSMUST0000003870) at 12 h after induction, which is the turning point from pre-BI to pre-BII stages. These 2 isoforms contain the helix-loop-helix DNA-binding domain but differ in their N-terminal sequences (Supplementary figure S2), Mxi1-202 being a longer protein. Interestingly, N-terminal variations of the human Mxi1 isoforms have been described to be associated with cytoplasmic retention of Mxi1 and fine control of *Myc* repression (Engstrom *et al*, 2004). The major isoform switch revealed by our *PodiumChange()* function may suggest that isoform-control of Mxi1 activity may also occur in murine B-cell differentiation.

## 4 Discussion

The Iso-maSigPro set of functions updates the maSigPro framework to analyze isoform changes in time course transcriptomics data. We model differential isoform utilisation as the interactions between the isoform, experimental condition and time, and evaluate significance with the log-likelihood ratio statistic of the models including or not this interaction. This approach selects genes where the relative proportions of their transcripts change in time. However, isoform expression differences might be small or only affect low expressed isoforms. To extract biologically meaningful changes in relative isoform abundances, we in troduced new clustering and querying functions. *seeDS()* and *tableDS()* help to find genes with substantial isoform profile differences in time, while *PodiumChange()* identifies those cases with a switch in the most expressed transcript. We showed examples where these functions helped to select genes with functionally relevant isoform changes. maSigPro is the first Bioconductor package with specific functions for the identification and analysis of alternative isoform expression in multiple time course transcriptomics experiments.

## 5 Acknowledgements

This work has been funded by the European Union Seventh Framework Programme STATe-gra project under the grand agreement 306000 and the Spanish MINECO grants BIO2012-40244 and BIO2015-71658-R. We thank Jesper Tegner and David Gómez-Cabrero for facilitating access to the RNA-seq data used in this study.

## References

AlShareef,S. et al. (2017) Herboxidiene triggers splicing repression and abiotic stress responses in plants. BMC Genomics, 18(1), 260.

Andres,S. et al. (2012) Detecting differential usage of exons from RNA-seq data. Genome Research, 22, 2008-2017.

Au,K.F. et al. (2013) Characterization of the human ESC transcriptome by hybrid sequencing. Proc. Natl. Acad. Sci. USA, 110, 50, E4821-30.

Conesa,A. et al. (2006) maSigPro: a Method to Identify Significantly Differential Expression Profiles in Time-Course Microarray Experiments. Bioinformatics, 22, 9, 1096-1102.

Engstrom,L.D. et al. (2004) Mxi1-0, an alternatively transcribed Mxi1 isoform, is overexpressed in glioblastoma. Neoplasia, 6, 5, 660-73.

Huang,Y., and Sanguinetti,G. (2016) Statistical modeling of isoform splicing dynamics from RNA-seq time series data. Bioinformatics, 32, 2965-72.

Jia,C. et al. (2016) MetaDiff: differential isoform expression analysis using random-effects meta-regression. BMC Bioinformatics, 16, 208.

Ma,S. et al. (2010) Ikaros and Aiolos inhibit pre-B-cell proliferation by directly suppressing c-Myc expression. Mol. Cell. Biol., 30, 17, 4149-58.

Matthew,E. et al. (2015) limma powers differential expression analyses for RNA-sequencing and microarray studies. Nucleic Acids Res., 47, e47.

McCullagh,P., and Nelder,J.A. (1989) Generalized linear models. Chapman & Hall/CRC, Boca Raton, Florida, 2nd edition.

Nueda,M.J. et al. (2014) Next maSigPro: updating maSigPro bioconductor package for RNA-seq time series. Bioinformatics, 30, 18, 2598-602.

Roberts,A., and Pachter,L. (2013) Streaming fragment assignment for real-time analysis of sequencing experiments. Nat. Methods, 10, 71-73.

Steijger,T. et al. (2013) Assessment of transcript reconstruction methods for RNA-seq. Nat. Methods, 10, 1177-1184.

Topa, H. and Honkela, A. (2016) Analysis of differential splicing suggests different modes of short-term splicing regulation. Bioinformatics, 32, i147âɘ§i155.

Vuong,C.K. et al. (2016) The neurogenetics of alternative splicing. Nat. Rev. Neurosci., 17, 265-81.

Wood,S.N. (2006) Generalized additive models. An introduction with R. Chapman & Hall/CRC.

